# The effect of different milk pretreatment methods on microbiome community development during Herrgårds cheese production and ripening

**DOI:** 10.1101/2025.04.23.648337

**Authors:** Juan Antonio Rodríguez, Luisa Santos-Bay, Apurva Narechania, Christian Carøe, Kimmo Sirén, Sarah S. T. Mak, Ida Broman Nielsen, Max Ramsøe, Thomas S. Pontén, Søren Lillevang, Lene Tranberg Andersen, M. Thomas P. Gilbert

**Author notes:** Equal contribution (Juan Antonio Rodríguez, Luisa Santos-Bay).

## Abstract

One of the biggest challenges for dairy producers is the substantial variability in final product properties caused by changes in the production environment. In cheese production, this variation is influenced by several factors, particularly the milk base and its pretreatment, which shape the microbiome throughout the process and ultimately affect the cheese’s organoleptic characteristics. To examine the impact of three different pre-treatments for pasteurised milk— microfiltration, protein fortification, and only pasteurisation (control)— on microbiome dynamics, we generated metagenome sequencing data from 14 cheese production steps across these three production trials at a Danish dairy factory. We constructed three metagenomic co-assemblies, identifying nine high-quality metagenome-assembled genomes (MAGs). Our analysis revealed that a specific strain of *Lactococcus lactis* dominates the process, while other minor bacterial species persist at very low abundances (<1%), contributing non-negligibly to product properties. Notably, *Clostridium tyrobutyricum*, a known dairy spoilage bacterium, was present at low levels in pasteurised-only and protein-fortified milk trials but was nearly absent in microfiltered milk. To enhance our analyses, we implemented KHILL, a novel *k*-mer-based method applied directly to raw sequencing reads, which facilitates metagenomic co-assembly and enables early detection of unwanted microorganisms. Our findings provide industrial dairy producers with a comprehensive view of microbial dynamics during cheese production, offering insights to improve process consistency and product quality.

## Introduction

Cheese has been produced through coagulation of the animal milk protein casein for at least 7,500 years (Salque et al. 2013). Over the last century its production has largely transitioned from an artisanal and manual cheesemaking process to a more industrialised one, linked to factories that are able to produce the volumes needed to fulfil the demands of its consumer base. However, a major challenge that industrial cheese factories often face, is the lack of product homogeneity when production environment is changed (Johnson, Curtin, and Waite-Cusic 2021). This is critical to industrial production, as even small changes in the process may have major consequences on the resulting organoleptic and rheologic properties of cheese. While there may be several explanations as to why this happens, it is almost certain that differences that arise in the environmental microbial communities during the manufacturing process play an important role in this variance (Fox et al., 2000.; Mayo et al. 2021). In cheese production, starter cultures that contain lactic fermenting microbes are added to the milk in a controlled and standardised way. However, other sources of microorganisms, like those naturally present in the milk or even those in the facility’s immediate indoors environment and the vicinity, may well enter the process and contribute to shaping the quality and properties of the final cheese. Factors that can alter the microbial communities include: the dairy where the cheese is made, the starter culture used, differences in equipment in the facilities, and even the sources of, and pre-treatment of, the milk.

Pre-treatment of milk is an increasingly common procedure in the dairy industry (Kelly, Huppertz, and Sheehan 2008). However, the effect of pre-treatment technologies, such as microfiltration and protein fortification is under-explored with regards to their effects on the cheese microbiome. In light of this, we initiated a study to profile how the microbiome community develops throughout a pilot plant trail mimicking an industrial continental cheese making process, as conditioned by the use of different starting milk bases and treatments, specifically microfiltration or protein fortification. Additionally, as the physical location in the cheese from where the sample is taken has been shown to bias downstream microbiome analyses (Irlinger and Monnet 2021), this was considered as a factor in our analyses.

Thanks to recent developments in culture-independent microbial community profiling through tools such as shotgun metagenomics (Warnecke et al. 2007; Stein et al. 1996), when adequate samples are available for analysis, it is now possible to reconstruct both the species within microbial communities, and the genes present in such microbes, at a level that was previously unachievable. Thus, in light of the need to better understand the factors that help shape variation in the industrial cheesemaking microbiome, we used this approach to the production of Herrgårds cheese at a single pilot plant facility in Skejby (Århus, Denmark), in order to explore how variation of key production parameters shaped the relationship of the microbiome and the final cheese. Herrgårds is a traditional Swedish hard cheese made from cow’s milk, with a pale yellow to golden-yellow interior with small, evenly distributed holes or “eyes” throughout the cheese. It has been described as being mild, nutty and creamy to taste (Jiang, Björck, and Fondén 1997). Specifically, we generated and compared the microbiome profiles for 3 pilot plant trials (PT1, PT2, PT3) of the cheese making process, taking samples at the same processing time points for all 3 trials (n = 14 samples *per* trial; total sampling points = 42). All trials started from the same source of pasteurised milk, and were subject to different ripening temperatures, samplings from different places in the cheese and different ripening times. In each of the 3 production trials (PT) the raw pasteurised milk underwent a different treatment, namely: microfiltration (PT1), only pasteurisation (PT2) and protein fortification (PT3). (Figure 1, see methods for details).

**Figure 1:**
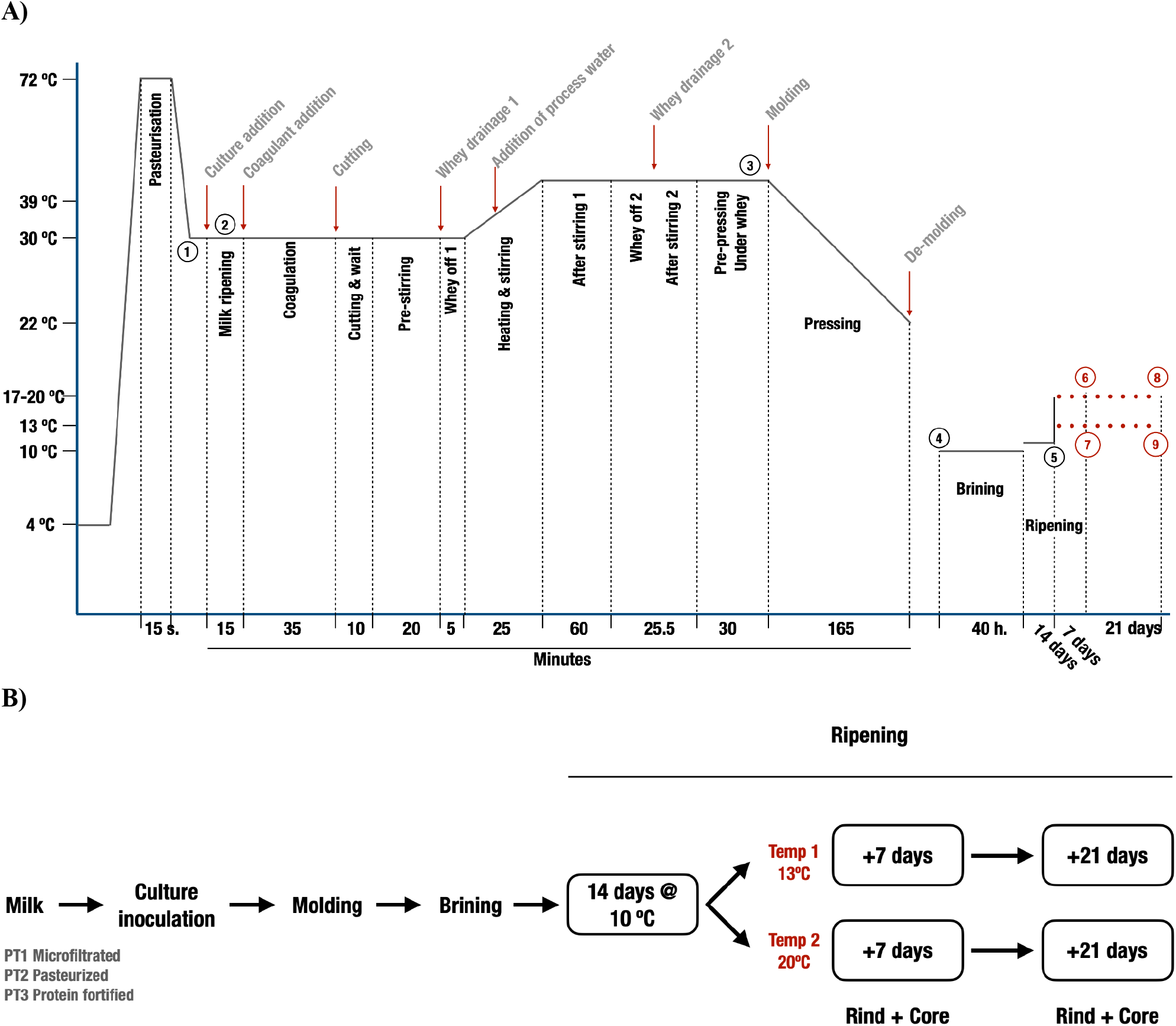
A) Diagram showing temperature and time throughout the cheese making process, indicating the sampling points where our samples were taken. B) Schematic diagram depicting the experimental setup for the trials. Sampling points n = 14.

Overall, our findings may help the dairy industry ensure robust production of premium quality cheeses, even if production is moved to different locations or different milk bases and pre-treatments are applied. This in turn will increase not only the cost-efficiency of cheese production in general, but also the overall cheese quality and in the end also benefiting the consumer.

## Results and Discussion

### Initial metagenomics shotgun data preprocessing

We generated shotgun metagenomic sequencing data from 14 different subsamples (Figure 1, Table 1) taken during each of 3 pilot trials of Herrgårdsost cheese, yielding 42 samples in total. Sampling timepoints included (Table 1, Figure 1A; following numbers in parentheses are referred to numbers in Figure A): (1) The source milk prior to inoculation with the industrial starter culture that is a blend of two *Lactococcus lactis* strains: subspp. *cremoris* and subspp. *lactis*, and one *Leuconostoc pseudomesenteroides* strains. Source milk was varied by production trial (PT) as follows: PT1: pasteurised + microfiltered, PT2: pasteurised only, and PT3: pasteurised + protein fortified, to explore how these processes may shape the subsequent community. Protein fortification is a commercial procedure in order to reduce to a minimum the defects such as precipitation, flocculation and sandiness of milk, due to the high heat tolerance of proteins (Lin et al. 2016). (2) Milk after culture inoculation, (3) before moulding and (4) before salt brining. Then (5 to 9), from both the cheese core and rind were sampled at multiple time points during the ripening phase, as described next.

**Table 1:**
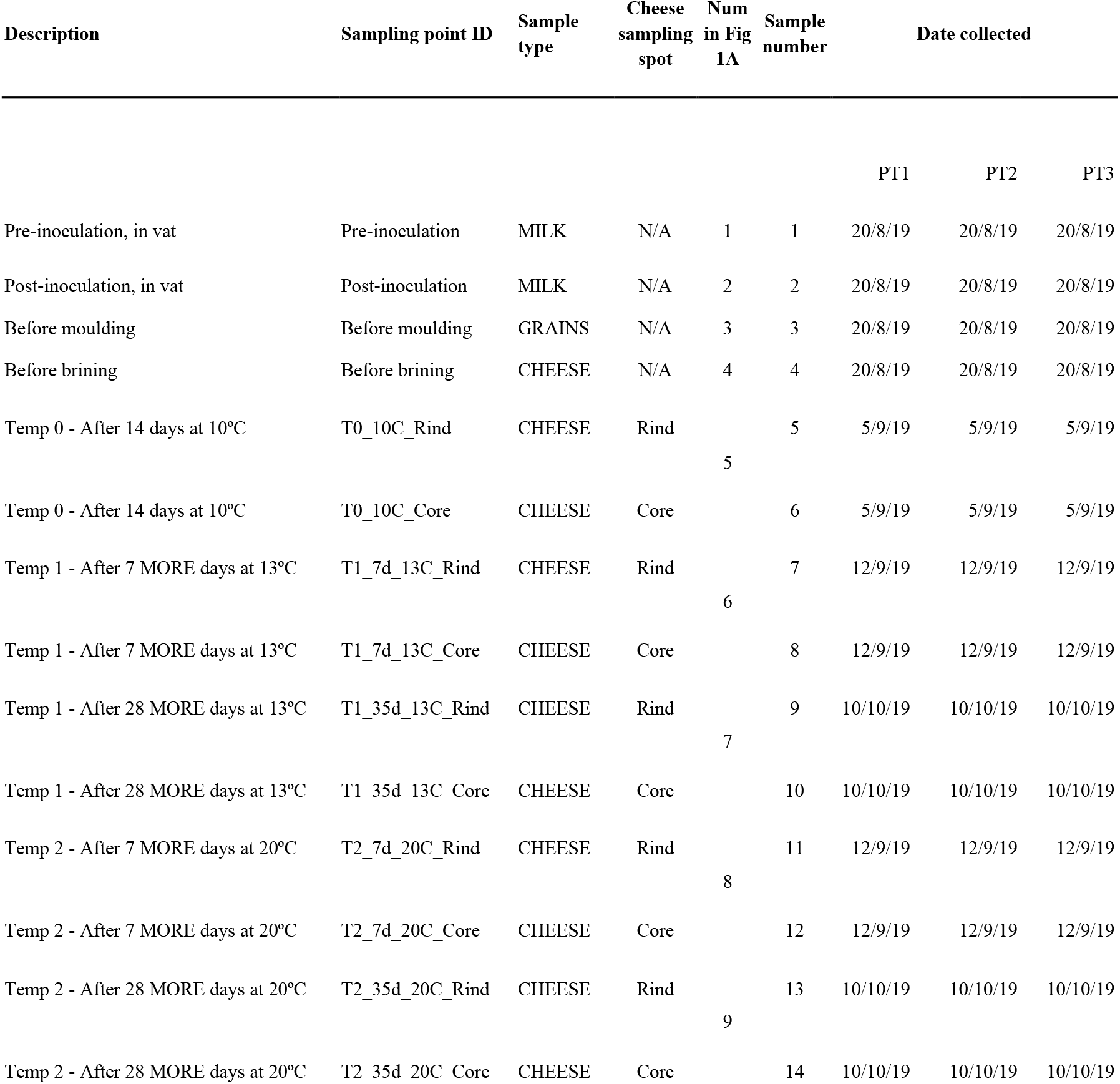
Description of samples and timepoints used. The column “Number in T^a^ graph” refers to the sampling points shown in Figure 1A.

| Description | Sampling point ID | Sample type | Cheese sampling spot | Num in Fig 1A | Sample number | Date collected |  |  |
| --- | --- | --- | --- | --- | --- | --- | --- | --- |
|  |  |  |  |  |  | PT1 | PT2 | PT3 |
| Pre-inoculation, in vat | Pre-inoculation | MILK | N/A | 1 | 1 | 20/8/19 | 20/8/19 | 20/8/19 |
| Post-inoculation, in vat | Post-inoculation | MILK | N/A | 2 | 2 | 20/8/19 | 20/8/19 | 20/8/19 |
| Before moulding | Before moulding | GRAINS | N/A | 3 | 3 | 20/8/19 | 20/8/19 | 20/8/19 |
| Before brining | Before brining | CHEESE | N/A | 4 | 4 | 20/8/19 | 20/8/19 | 20/8/19 |
| Temp 0 - After 14 days at 10°C | T0_10C_Rind | CHEESE | Rind | 5 | 5 | 5/9/19 | 5/9/19 | 5/9/19 |
| Temp 0 - After 14 days at 10°C | T0_10C_Core | CHEESE | Core |  | 6 | 5/9/19 | 5/9/19 | 5/9/19 |
| Temp 1 - After 7 MORE days at 13°C | T1_7d_13C_Rind | CHEESE | Rind | 6 | 7 | 12/9/19 | 12/9/19 | 12/9/19 |
| Temp 1 - After 7 MORE days at 13°C | T1_7d_13C_Core | CHEESE | Core |  | 8 | 12/9/19 | 12/9/19 | 12/9/19 |
| Temp 1 - After 28 MORE days at 13°C | T1_35d_13C_Rind | CHEESE | Rind | 7 | 9 | 10/10/19 | 10/10/19 | 10/10/19 |
| Temp 1 - After 28 MORE days at 13°C | T1_35d_13C_Core | CHEESE | Core |  | 10 | 10/10/19 | 10/10/19 | 10/10/19 |
| Temp 2 - After 7 MORE days at 20°C | T2_7d_20C_Rind | CHEESE | Rind | 8 | 11 | 12/9/19 | 12/9/19 | 12/9/19 |
| Temp 2 - After 7 MORE days at 20°C | T2_7d_20C_Core | CHEESE | Core |  | 12 | 12/9/19 | 12/9/19 | 12/9/19 |
| Temp 2 - After 28 MORE days at 20°C | T2_35d_20C_Rind | CHEESE | Rind | 9 | 13 | 10/10/19 | 10/10/19 | 10/10/19 |
| Temp 2 - After 28 MORE days at 20°C | T2_35d_20C_Core | CHEESE | Core |  | 14 | 10/10/19 | 10/10/19 | 10/10/19 |

The steps of the production were then standardised up to the ripening stage, where after an initial 14 days at 10º C (T0), each batch was subdivided into one ripening at 13º C (T1) and a second at 20º C (T2), in order to explore if temperature variation played a significant role on the microbiome community (Figure 1B). Post DNA extraction, shotgun microbiomes were sequenced using 2×100 bp PE Illumina sequencing, with an average output of 52.5 M (s.d: 15.2 M) pairs of reads generated for each of the 42 libraries. Reads were trimmed to remove sequencing adapters, leaving an average of 99.43% of the initial raw reads (Table S1). Afterwards, endogenous host DNA (cattle; *Bos taurus*) was removed by mapping against the cow reference genome (ARS-UCD1.3, *bosTau9*). Up to 99% of the reads were removed in early raw milk stages, while removing <1% during ripening (Table S1).

### k-mer analyses reveal stable composition of cheese microbiome

A critical first step in the analysis of shotgun metagenome data is the co-assembly of sequencing data generated from each independent sample, into a resulting metagenome-assembled genome (MAG) catalogue, against which the taxonomic composition profiling of each sample can eventually be obtained. This process in turn requires a prior decision to be made about which data should be pooled for each coassembly, as the computational efficiency of the process depends on balancing inclusion of too many samples with too diverse microbiomes, against inclusion of too few samples - both can give rise to a poorly assembled MAG catalogue. To inform this decision, we first analyzed the raw FASTQ shotgun sequence data using KHILL, a novel k-mer counting method based on Hill numbers (Narechania et al. 2022). This approach enables computationally efficient exploration of sequence diversity directly from metagenomes, without requiring assembly or taxonomic profiling, even at shallow sequencing coverage. Fundamentally, KHILL is an information diversity-based technique that calculates the effective number of metagenomic samples given the distribution of *k*-mers between those samples. Completely unique, non-overlapping samples with no information in common would result in a KHILL value of 2 (two effective samples). Any *k*-mer overlap would lower the KHILL statistic. For pairwise comparisons, the metric therefore varies between 1 and 2. A KHILL of 1 indicates completely identical samples in terms of both *k*-mer identity and frequency. Initially designed to rapidly detect community shifts from a baseline, the approach has been previously used to efficiently detect temporal variation in SARS-CoV-2 viral strains sequenced from wastewater collected during the 2020 pandemic period (Narechania et al. 2022). This tool has two benefits in the context of our study, relating to the rapid insights it provides into the similarity/dissimilarity of the different samples. Firstly it provides a rapid overview of the stages at which the community is changing (something that ultimately may represent an attractive tool for cheese producers who may wish to dynamically profile the microbiome community in their samples), and secondly these observations can be used to guide the co-assembly binning decisions.

We therefore ran KHILL (Narechania et al. 2022) for each of the 14 timepoints for the 3 PTs (Figure 1). We used 3 million reads randomly selected from each of the 14 sampling points in each PT as an input for KHILL and obtained correlation matrices shown in Figure 2. For all 3 batches we see similar patterns, with the ripening phases after moulding (T0, T1, T2) being quite stable, with no major compositional changes.

**Figure 2:**
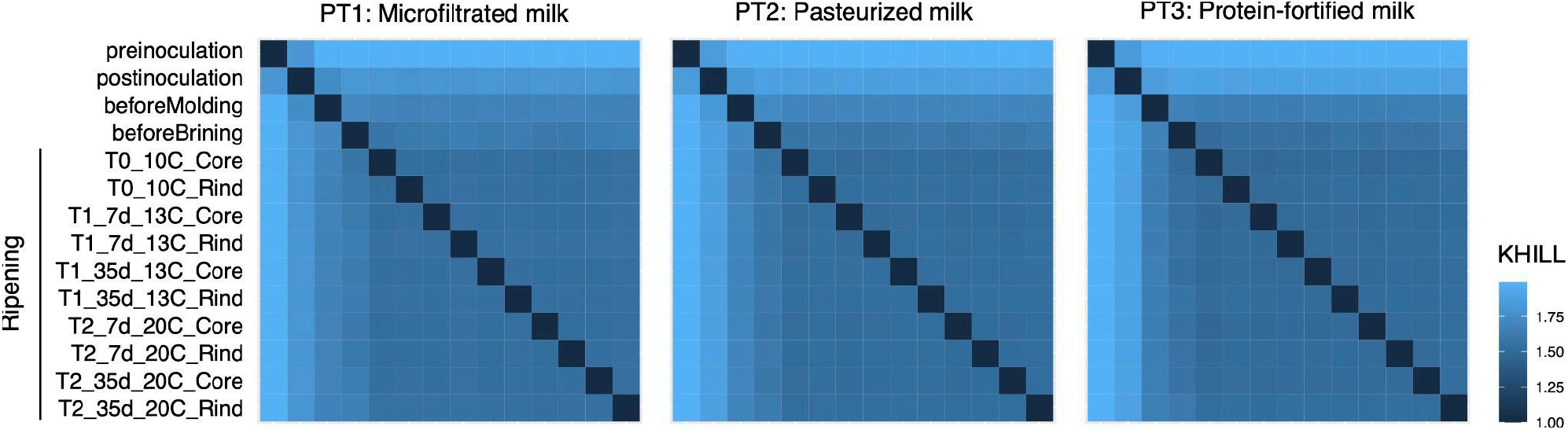
Heatmaps showing the similarity or differences in k-mer composition between the samples. The scale colour represents the intensity of the differences in k-hill observations. Darker intensities denote smaller differences in composition.

In all PTs, principal differences observed were between the ripening phase (steps T0, T1, T2) and the first phases (“Pre-inoculation” and “Post-inoculation”; liquid milk sample), and “Before moulding” and “Before brining” (cheese grains sample). This can be attributed to fluctuations in microbial composition occurring at key processing stages. Specifically, we observe compositional shifts from raw milk (pre-inoculation) to pasteurised milk following the addition of starter cultures (post-inoculation). As fermentation progresses, further changes occur, culminating in the microbial profile observed in coagulated cheese grains (before moulding). KHILL comparisons between pre-ripening stages, and between these stages and ripening, reflect wholesale changes in the microbial composition in these early stages, and starter culture expansion in the cheese (KHILL ∼ 2). Finally, we see the next shift just before entering the ripening process (Before brining). Note that all these changes happen in less than 10 hours, while the ripening process lasts for up to 35 days, which in this particular case informs us about the stability of the microbial composition along the ripening.

The above observations allowed us to form two conclusions. Firstly, the stability and low compositional changes at the ripening steps for all 3 PTs prompted us to generate a metagenomic coassembly independently for each of the replicates in order to validate that, indeed, the composition is stable. Secondly, given that an estimated ∼10-20% of milk and dairy production worldwide is lost due to bacterial spoilage (Dousset, Jaffrès, and Zagorec 2016) and thus there is an obvious interest in both closely controlling the production process, and being able to rapidly detect microbial changes that may affect the sample quality (Ziyaina, Rasco, and Sablani 2020), KHILL may represent an interesting tool for integration into the cheese-making process. Specifically, as a result of the decreasing cost and increasing outputs of novel high-throughput sequencing technologies such as those marketed by Oxford Nanopore Technologies (ONT), they are beginning to be applied for real-time monitoring of environmental and industrial bioprocesses (Kovaka et al. 2021; Bruno, Aury, and Engelen 2021). Unfortunately, however, the data generated conventionally requires considerable subsequent computational processing such as mapping reads to a database of choice. This is a time-costly bottleneck step that could be resolved by the application of the less computationally demanding KHILL method.

### MAG diversity quantitatively confirms stability across cheese production trials

We generated 3 metagenomic co-assemblies from the data (Methods) using the Anvi’o workflow for metagenomics (Köster and Rahmann 2012; Eren et al. 2021), one for each of the 3 PTs, involving all 14 samples per PT (Methods, Table S1). Using the Anvi’o interactive interface we manually binned the data into 5 (PT1), 9 (PT2) and 10 (PT3) metagenomic bins, with a minimum of 36% observed completion, as estimated by Anvi’o collection of single-copy core genes (SCG) (Methods) (Table 2). After the co-assembly, we obtained 411, 3,025 and 3,391 contigs respectively for the 3 PTs, with a length equal to, or longer than, 1 Kb (Table 2). We retrieved a total of 18 different bins belonging to the *Bacteria* taxonomic domain (Table S2) for the 3 PTs, plus 6 bacteriophage-containing bins (discussed below). Taxonomy up to the level of species was assigned by Anvi’o program anvi-run-scg-taxonomy, using Anvi’o SCG collections (Lee 2019), while the interactive manual binning was in process. Additionally, contig-level taxonomic classification was determined by Kaiju (Menzel, Ng, and Krogh 2016), and incorporated into the Anvi’o interactive visualisation.

**Table 2:** Basic statistics from the coassembly process for each of the pilot trials.

| ID | Total contigs | Longest contig | Estimated number of genes | Total contig length | N50 | Anvi'o estimated bacterial genomes | Total selected bacterial bins | Total contigs assembled into bins | % |
| --- | --- | --- | --- | --- | --- | --- | --- | --- | --- |
| PT1 | 2952 | 137 kb | 14207 | 11.97 Mb | 10.18 kb | 3 | 3 | 411 | 13.92 |
| PT2 | 5163 | 136.8 kb | 23228 | 20.01 Mb | 7 kb | 6 | 7 | 3025 | 58.59 |
| PT3 | 5129 | 136.8 kb | 24910 | 21.62 Mb | 8.4 kb | 7 | 8 | 3391 | 66.11 |

From these 18 bacterial bins we generated a final unique MAG catalogue, consisting of a total of 7 high-quality unique bacterial MAGs (Table 3). For that, we manually dereplicated the 18 bins from the 3 PTs (Table S2), by keeping those bins with >70% genome completion and <5% redundancy. In case of a tie, the longest genome was kept. Of note, amongst all the MAGs included in the final catalogue, only the *Janthinobacterium spp*. MAG from PT2, was under 95% in completion (completion = 71%). For both *Lactococcus* subspecies MAGs, their corresponding starter culture representative genomes were selected as representative MAGs, after phylogenomic analyses identified the strain from the cultures present in our cheese (see Supplementary Information).

**Table 3:** De-replicated MAGs and statistics.

| MAG name | Origin | Length (bp) | Contigs | N50 | GC content | Completion (%) | Redundancy (%) | Species name |
| --- | --- | --- | --- | --- | --- | --- | --- | --- |
| CRE6 | CULTURES | 2510276 | 210 | NA | NA | 100.00 | 0.00 | *Lactococcus lactis cremoris* |
| LAC2 | CULTURES | 2421935 | 152 | NA | NA | 98.59 | 0.00 | *Lactococcus lactis lactis* |
| Leuconostoc_pseudomesenteroides | PT2 | 2001676 | 139 | 33436 | 37,45 | 100,00 | 1,41 | *Leuconostoc pseudomesenteroides* |
| Clostridium_tyrobutyricum | PT3 | 2630280 | 627 | 5530 | 30,49 | 97,18 | 2,82 | *Clostridium tyrobutyricum* |
| Janthinobacterium_spp | PT2 | 2567761 | 744 | 4550 | 62,19 | 71,83 | 2,82 | *Janthinobacterium spp.* |
| Lactacaseibacillus_paracasei | PT3 | 2869405 | 357 | 15310 | 44,87 | 98,59 | 0,00 | *Lactacaseibacillus paracasei* |
| Streptococcus_spp | PT3 | 1649953 | 200 | 15109 | 38,76 | 98,59 | 0,00 | *Streptococcus spp* |
| Ceduovirus_1 | PT2 | 22165 | 9 | 2851 | 35,26 | 100.00 | NA | *Lactococcus phage BIM BV-114, (predicted)* |
| Ceduovirus_2 | PT2 | 20592 | 6 | 4768 | 35,20 | 93,93 | NA | *Lactococcus phage bIL67, (predicted)* |

We obtained the lowest number of bacterial MAGs (n = 3) from the microfiltered milk trial (PT1), with no taxa found to be exclusively present in this trial (Table 2, Table S3). This low number matches the expectation of the effect of the microfiltration process (see Methods section), by removing any potential opportunistic taxa in milk. Specifically, these 3 MAGs match the 3 expected bacteria from the added starter cultures, namely: the two subspecies of *Lactococcus lactis*: subspp. *cremoris* and *lactis* (see Supplementary Information), plus one strain of *Leuconostoc pseudomesenteroides*, although its abundance is <1% out of all the reads recruited by the MAG catalogue.

In trials PT2 and PT3, both derived from milk that had not undergone microfiltration, we successfully assembled 7 and 8 MAGs, respectively. Of these, 7 were shared between both trials, including the 3 expected starter culture bacteria mentioned earlier. Following the MAG dereplication step, 4 unique MAGs detected in both PT2 and PT3 were retained in the final catalogue, along with the three from starter cultures, resulting in a total of 7 final bacterial MAGs. The 4 new MAGs identified in trials PT2 and PT3 were: *Lacticaseibacillus paracasei, Janthinobacterium spp*., *Clostridium tyrobutyricum* and *Streptococcus spp*. Genus *Janthinobacterium* is known to cause milk spoilage (Eneroth, Ahrné, and Molin 2000), while *Clostridium tyrobutyricum* confers a particularly undesired rancid smell and taste to cheese and can create late blowing (Dousset, Jaffrès, and Zagorec 2016). *Clostridium* is also known as one of the causative agents for late blowing defects observed in semi-hard and hard cheeses (Silvetti, Morand and Brasca, 2018), which can render the cheese not suitable for commercialisation. *Lacticaseibacillus paracasei* is considered a safe microorganism, naturally occurring through dairy processing (Falfán-Cortés et al. 2022). *Streptococcus* is a milk associated bacteria, which can be part of starter cultures, and is considered non-pathogenic (Yerlikaya and Ozer 2014; Dan et al. 2018). While our results indicate that these bacteria are found at relatively low levels (<1%) throughout the process, we observed that their abundances consistently increase after inoculation across PTs, and remain stable during ripening (Figure S1). All in all, despite their relatively low abundance, gene-expression and metabolic analyses would be needed in order to understand their true contribution towards the final properties of cheese. One additional bin, classified as *Pseudomonas spp*., was recovered from both PT2 (completion 36.6%) and PT3 (completion 60.6%). Despite the low completion and not satisfying the threshold for being a high-quality MAG, we still believe that it is worth reporting, as it confirms the low abundance of this potentially detrimental microorganism (Boor and Fromm 2006). And most importantly, it remained undetected in the microfiltered milk trial PT1, which again demonstrates the effectiveness of milk microfiltration (Figure S1).

### Viral genomes

In addition to the 7 bacterial MAGs, we were able to recover from each of the 3 PTs, 2 further bins of non-bacterial origin, with their contigs classified by Kaiju (Menzel, Ng, and Krogh 2016) as *Lactococcus* bacteriophage viruses (Family *Siphoviridae*). Their average genome length was 19.4 kb (min: 12.9 kb; max: 23.2 kb) (Figure 3, Tables S3-5), congruent with the size of a complete phage genome (Deveau et al. 2006). Initially, before deduplication and selecting them as MAGs, we named the bins *Ceduovirus_1* and *Ceduovirus_2*, as Kaiju classified one of the bins with this name, while the other bin was assigned to the category of *Unknown_Siphoviridae*. The *Ceduovirus* genus (alternatively known as “*c2*” virus) is one of the 3 most common groups of lactococcal phages, with a genome size of ∼22 kb (Hulo et al. 2011), which fits our results (Table S5), with the other 2 most common groups (named 936 [or *Skunavirus*] and P335 phages, respectively), both having larger genomes than *Ceduovirus* (Deveau et al. 2006).

**Figure 3:**
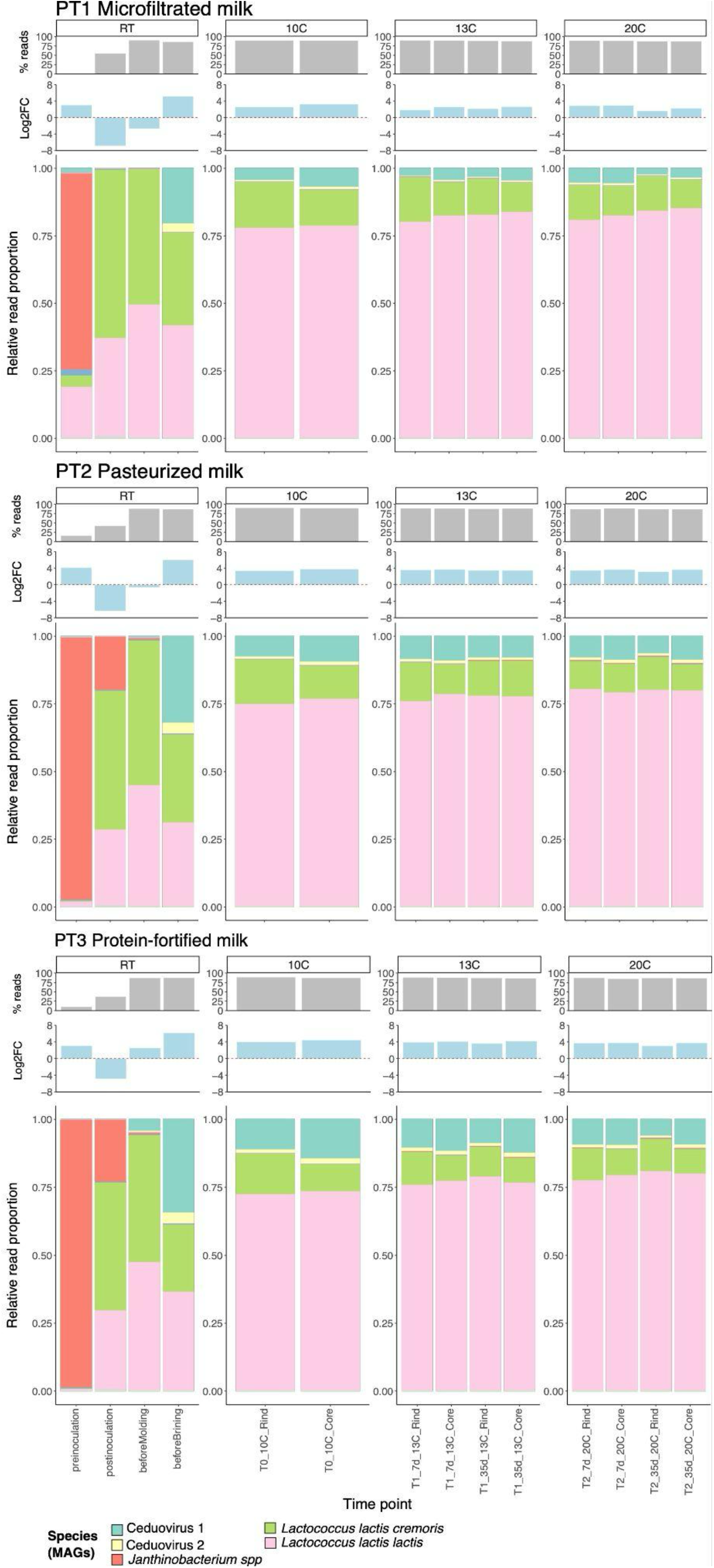
Relative abundance estimated as % of reads mapping to the 9 MAGs generated, for each of the 3 replicates. **Top panes:** percentage of reads from the raw data recruited by the MAGs. The “preinoculation” step recruits a low percentage of reads because of the initial low amount of microbe raw data at that step after host removal, as described earlier. **Middle panes:** log2 ratio for the proportion of phages to bacteria, estimated through coverage ratios with log-fold change applied. **Bottom panes:** Relative abundance for the 9 MAGs, in percentage of reads recruited.

Anvi’o is not designed to work with viral genomes. Thus, in order to estimate the completeness of the phage genomes we used CheckV, v.0.9 (db version 1.2) (Nayfach et al. 2021) (Methods). CheckV returned that completion for the bins was >93% for 4 out of the 6 phage genomes detected (2 per PT trial), with the remaining 2 phages being assigned as 58% and 73% complete (Table S4). As we do not know the full taxonomic classification for the phages, and as both of them appear closely related, we derreplicated the genomes using the Anvi’o command anvi-dereplicate-genomes, which relies on computing average nucleotide identity (ANI) between all 6 genomic phage bins detected. Anvi’o determined that with at least 90%, and no more than 95% ANI, there were two different clusters. Anvi’o assigned exactly one bin from each PT to each phage group, fitting our hypothesis of having two distinct phage strains. We selected the two most complete genomes as representatives of each phage group, both obtained from PT2. These genomes had completion rates of 100% (22.2 kb) and 94% (20.6 kb), respectively, and were classified as “high-quality” by CheckV. They were subsequently added to our MAG catalogue, bringing the total to 9 unique species (Table 3). In an attempt to further determine the exact taxonomic identification of the phages, we ran BLAST v.2.2.5 (Altschul et al. 1990) with our viral MAGs against the NCBI nt nucleotide database (accessed 22/08/2022), and we ranked results by the identity (>92%), %query coverage and length of the match (Tables S7-8). Top hits revealed two lactococcal phages, namely phage *BIM BV-114* and phage *bIL67* (Table 3).

### A relatively low number of MAGs

In light of our observations, an obvious question is whether the recovery of relatively few MAGs is expected, or rather reflects a limitation of our use of a shotgun sequencing approach. In this regard we note that prior studies of the cheesemaking process based on metabarcoding based analyses of 16S diversity report similar magnitudes of OTUs. For example, in one recent 16S based study, a maximum of 10 identified species were recovered from the surfaces of Époisses cheeses, both across ripening (28 days) and storage (90 days) (Irlinger and Monnet 2021). A second study that profiled the microbiome of industrial cheese using 16S metabarcoding identified a median of 6.5 amplicon sequence variants per cheese sample (Johnson, Curtin, and Waite-Cusic 2021), thus fitting our observations. In a third study that examined the microbial communities of washed-rind cheese produced from raw and pasteurised milk in an artisanal, non-industrially controlled process, researchers identified between 11 and 37 microbial species from the cheese rind (Saak et al. 2023). Our results fall within the lower range of this analysis, which may be attributed to the shorter ripening times in our study. We did not detect a significant level of fungi in our reads (<0.01% of all the reads in PT3, as determined by Kaiju (Menzel, Ng, and Krogh 2016). Several studies indicate that bacteria generally outnumber fungi by orders of magnitude through cheese production (Quijada et al. 2020; Irlinger and Monnet 2021), even in cheese that have been only salt brined, as it is our case.

### Lactococcus spp. strains stably dominate throughout the ripening process

After assembling a unique MAG catalogue, we sought to quantify the abundance of each of the 9 microbial species across the 14 key stages of the cheese production process. To do this, we mapped back our host-depleted raw metagenomic reads against the MAG catalogue, and quantified the relative abundance as percentage of reads recruited by each MAG. Broadly, for all PTs, we observed a relatively stable and constant trajectory for the abundances of MAGs, with principal domination by *Lactococcus*, as previously reported (Johnson, Curtin, and Waite-Cusic 2021) (Figure 1, 3). This aligns well with our previous analysis using *k*-mers, where we observed a high compositional stability through ripening time (Figure 2). As expected, at the pre-inoculation step only a small fraction of all the reads were recruited to the MAG catalogue (PT1: 0.6%, PT2: 16%, PT3: 9.9%) (Figure 3). PT1 has a remarkably lower amount % of reads mapping back to the MAGs at this step, as this batch was additionally microfiltered, theoretically removing any other milk microorganisms, despite the raw number of reads before being of comparable magnitudes: PT1: 244k, PT2: 306k, PT3: 177k (*in read pairs*). Immediately after the inoculation step, the percentage of reads mapped increased to ca. 40-50% for all 3 batches, with *Lactococcus* strains broadly dominating all the batches (Figure 3). At this point, for all 3 trials, the proportion of *cremoris* over *lactis* subspecies had a -log2FC of ∼ 1 (Figure S2), meaning that the proportions of both are equally balanced. The trend is then inverted after the salt brining bath and *lactis* starts outnumbering *cremoris*, until stabilising at between 2-3x -log2FC, specially through the ripening process. The metabolic capabilities of ssp. *lactis* (Supplementary Information) may help to explain this differential thriving, as has been also previously reported (Fernández et al. 2011). The overall domination of lactococcal bacteria has also been recently reported in (Irlinger and Monnet 2021), with more than 70% of relative abundance for two *Lactococcus* subspecies *(L*.*l. lactis* and *L. l chungangensis)* through the ripening stages (28 days ripening). Incidentally, subsp. *lactis* seems also to be here the dominant one across the whole process.

Finally, for almost all of our time points beyond the “Before moulding” step, 85 to >90% (Figure 3) of the reads are recruited to our MAG catalogue, which illustrates the fact that our MAG collection captures quite well the microbial communities in our pilot plant cheese samples.

### Bacteriophage viruses are ubiquitous through the cheesemaking process

Throughout the whole process, we detected a considerable abundance of the 2 lactococcal phage MAGs (Figure 3). Phages are known to disrupt starter cultures, particularly *Lactococcus* phages (Spus et al. 2015). Notably, at least one phage strain was already present in the milk, with mean coverage values below 1X across all PTs, including microfiltered milk. This suggests that viable phages can persist despite both pasteurization and microfiltration, albeit in reduced quantities (Garneau and Moineau 2011). To obtain an estimate of the proportion of bacterial cells to phage virus, we computed the relation between their genome coverages, and turned it into a log2 fold change (log2FC). Immediately after the addition of starter cultures, most of the sequences generated consisted of *Lactococcus*, and the ratio of phage units to *Lactococcus* cells became negative, for all 3 PT (more bacteria than virus). It is not until the brining bath step that this trend is inverted, reaching a log2FC >4, for all 3 PTs (Figure 3, middle panel). Brining salt baths are known to reduce the microbial complexity of cheese (Mounier et al. 2017), thus this may have helped to slightly reduce the amount of phages, before entering the ripening stage. Then, once in the ripening process we observed a remarkably continuous and consistent ratio between phages and *Lactococcus* within batches. After the brining bath, and once cheeses enter the ripening process, the microfiltered milk PT1 showed a consistently lower proportion of phages to *Lactococcus* cells, with an average log2FC of 2.44, while for PT2 and PT3 it was 3.5 and 3.83, respectively (Figure. 3). Unfortunately, we do not have replicates for the microfiltered milk PT, and thus we have no statistical power to demonstrate that microfiltration consistently reduces phage content, but we are confident this observation is worth to report, as this relationship phage:bacteria was stable alongside the whole ripening process. We can conclude that microfiltration is not able to remove all phages in the milk, but our limits in experimental design do not allow us to prove whether they come from inside the factory infrastructures.

Both phage strains seem to have one particular *Lactococcus* strain as their target host. The proportion of abundances between the 2 bacterial hosts and the 2 viruses seem to correlate quite well in general terms and the correlation coefficient between proportions of either lactococcal strains and their respective phages are ∼90%, while also significant (pv << 0.05) (Figure S3). We interpret this as that there is not an imbalance regarding the growth of phages or bacteria, which keep their respective growths linked, being remarkably significant during the ripening phase.

### Cheese rinds and cores showed no lactococcal community differentiation

We quantified how different the rind and core communities of the cheese were in terms of *Lactococcus* communities. For our Herrgårds cheese examined here, *Lactococcus* is the main bacterial group involved in the cheese final characteristics, so the sum of both *Lactococcus* strains’ abundances was used as a response variable in a two-way ANOVA analysis. ANOVA was run to determine whether the cheese sampling point (rind or core) has any effect on the abundance of *Lactococcus* recovered, and if this is different across trials. Through the ripening process (steps 5-9 in figure 1) a total of 15 data points were collected per each sampling site (5 per trial) and the average *Lactococcus* abundances per group were calculated (Table S9).

ANOVA assumptions were satisfied, as data did not deviate from normal distribution (Shapiro-Wilks test pv = 0.99) and variances between groups were not significantly different (Levene test pv = 0.27). An initial visual inspection of the data revealed that, in general, the rind tends to be more rich in *Lactococcus* than the core (Figure S4). ANOVA further demonstrated that there is both a significant effect (pv = 3.27e-09), when looking at the PT influence alone (*i*.*e*: the type of milk) in the amount of *Lactococcus*, but also the abundance is different between the sampling sites (pv = 2.93e-03). Interaction of site:trial was not significant (pv = 0.59), meaning that the change in abundance is not particularly significant after the combined effect of both factors. Our analysis had a statistical power ∼ 100% to detect an effect size as the observed for the PT factor at pv ≤ 0.05 (Effect size: Ω^2^ for PT factor = 0.71; site factor = 0.07), but unfortunately our design does not have enough statistical power (∼10%) to detect such a small effect size from the cheese sampling point. Despite not being powered enough, the abundance of *Lactococcus* was systematically higher for rind samples and, if more samples become available, the possibility remains open to statistically validate our hypothesis and observations.

### Less abundant bacteria may also leave a big imprint

Besides the landscape being dominated by *Lactococcus* strains, the remaining bacteria from our MAG catalogue are also present during the process. We recalculated relative abundances excluding both *Lactococcus* and viral MAGs (who dominate the dataset), and plotted percentages to better visualise the contribution of these 5 other minorly present bacteria, which have been also assembled into MAGs. On average, these minor MAGs represent less than 1% of the total composition of MAGs during ripening (Figure S1, Table S10). Right after the culture mix addition, *Leuconostoc pseudomesenteroides* becomes increasingly abundant and maintains a fairly constant abundance in all PT, even after 35 days of ripening. *Clostridium tyrobutyricum* is a bacteria that can be present in raw milk (Podrzaj et al. 2020). It shows up during later ripening stages conferring particular smell and taste to the cheese, which may not always be suitable for consumption and commercial production. We compared the abundance of *C. tyrobutyricum* at our PTs. First, we saw a highly significant reduction of *C. tyrobutyricum* in microfiltered trial PT1 compared to PT2 and PT3 (average abundance ratio: 4768x more abundant in PT2 and 4721x in PT3 than PT1. Both comparisons were significant: Two-tailed Wilcoxon test p-value: PT1-PT2 = 6.2e-4 | PT1-PT3 = 7e-3) (Figure 4B). Thus it seems clear that microfiltration removes nearly all *C. tyrobutyricum* present in the milk, which only starts showing up in PT2 and PT3 after 21 days of ripening (Figure S1).

**Figure 4:**
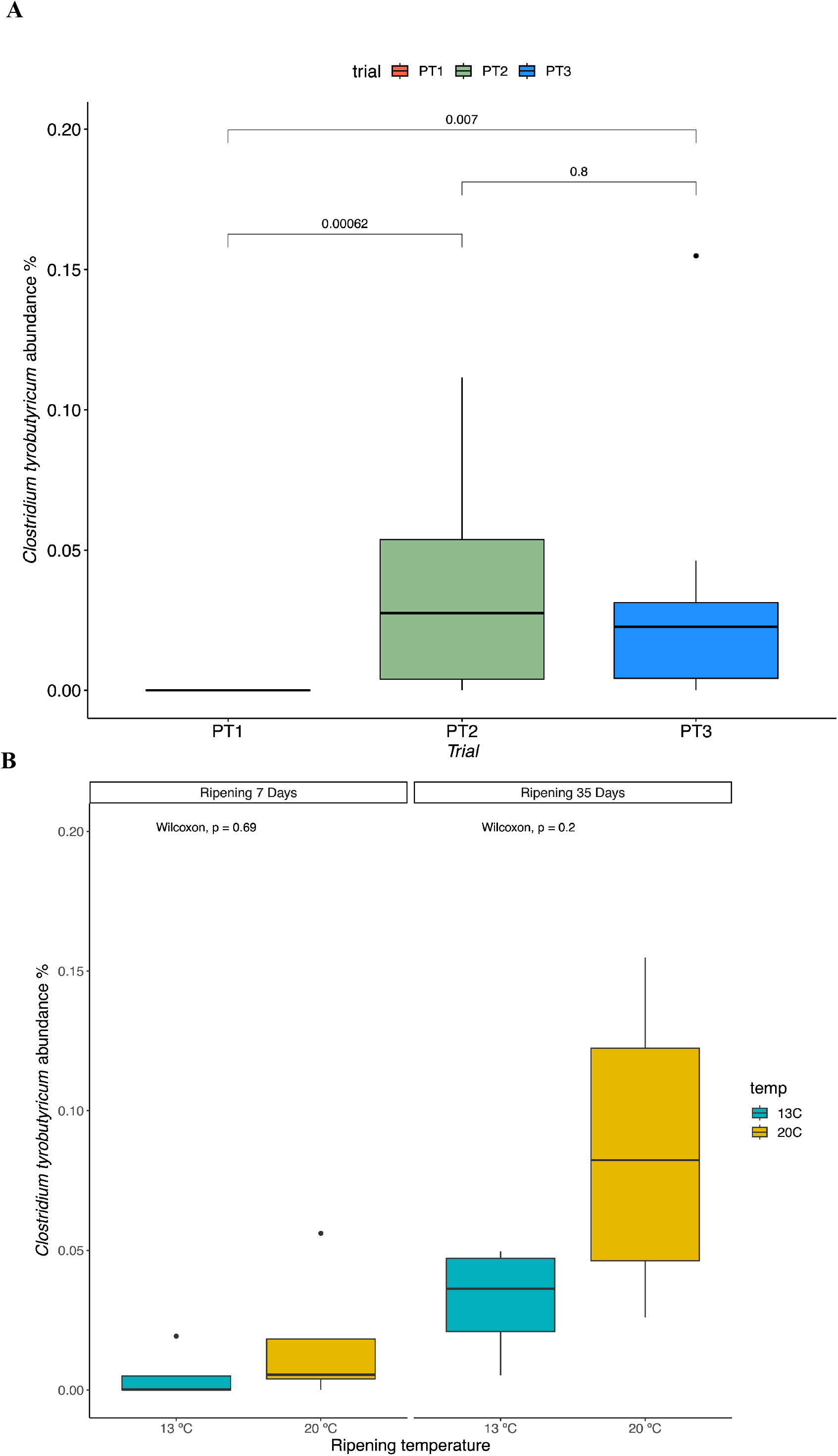
**A)** Total % abundance of C. tyrobutyricum at each trial. Numbers in the plot connecting groups denote p-values derived from pairwise Wilcoxon-tests between trials. Y-axis represents % abundance of C. tyrobutyricum across the different PTs. **B)** The effect of ripening days and temperature on C. tyrobutyricum growth (% abundance). Numbers in the plot denote p-values derived from pairwise Wilcoxon-tests within ripening days categories.

The following analyses do not consider PT1, as we can confidently say that the amount of C. tyrobutyricum cells is technically zero. We were interested in understanding C. tyrobutyricum growth patterns in relation to temperature and ripening days. Despite only having 4 data points at each temperature per number of ripening days, we found a clear trend (Figure 4B) showing that the high temperature set (20 ºC) promotes development of C. tyrobutyricum. This is confirmed by the fact that longer ripening times (35 days) show higher C. tyrobutyricum abundance than shorter ripening times (7 days ripening). At each group (temperature/days) shown in figure 4B we have only 4 data points, so we lack statistical power to perform proper comparison within ripening days, despite the trend towards higher temperatures being remarkable.

However, after grouping *C. tyrobutyricum* abundances by ripening day, independent of the temperature factor, ANOVA tests showed significant differences (p-value = 0.028), while temperature as a factor is not significant (p-value = 0.13). *C. tyrobutyricum* reads represent a very little percentage of the reads per each trial during ripening (PT1: 0.0006%; PT2: 0.281%; PT3 0.28%, Table S10), but despite this minimal abundance, organoleptic properties of the cheese might appear affected by this microorganism.

Despite the stability of the cheese microbiome along the process, the organoleptic properties of final cheese products were slightly different. This suggests that other low-abundance bacteria, either coming from our MAGs or not, can be active with their metabolic volatile compounds influencing the final characteristics of cheese. In order to quantify that, transcriptomics and volatile compounds analysis would be necessary to gain a full understanding of a process which, *a priori* and compositionally speaking, seems quite stable.

### Concluding remarks and perspectives

Recent advances in shotgun metagenomic sequencing and computational approaches have significantly enhanced our understanding of genome-resolved environmental metagenomics (Pinto and Bhatt 2024). In the food industry, these techniques have revolutionised our knowledge of microbial dynamics and their role in food production (Billington, Kingsbury, and Rivas 2022). Combined with the increasing accessibility of multi-omics data, they provide detailed insights into microbiome dynamics. Despite previous studies describing changes in the cheese microbiome during production (Irlinger and Monnet 2021; Johnson, Curtin, and Waite-Cusic 2021; Saak et al. 2023), a systematic evaluation of how milk pre-treatment affects microbiome evolution remains lacking. Our analysis of microbial composition during industrial cheese production reveals high stability and low diversity across the entire process, largely independent of milk pre-treatment. This low diversity is primarily driven by the stable dominance of two distinct *Lactococcus* strains and their associated phages during ripening. Notably, microfiltration effectively eliminated undesirable bacteria (e.g., *Clostridium* and *Pseudomonas*), reinforcing its prophylactic role by ensuring that only the intentionally introduced culture microorganisms remain. Although present in low abundance, other bacterial species exhibited a consistent presence across all three production trials. To fully understand microbial dynamics— including both dominant and low-abundance bacteria—future studies should integrate metagenomics (DNA), metatranscriptomics (RNA), and metabolomics to achieve higher-resolution insights into how milk pre-treatment affects the organoleptic properties of cheese. Our metagenomic analysis provides a valuable baseline framework, particularly for dairy producers interested in evaluating the effects of different milk bases. Additionally, our KHILL method, applied here to guide co-assembly, offers a practical application for the dairy industry. Easily implementable in a factory setting, KHILL can rapidly assess batch quality and detect potential spoilage early in the production process, allowing for timely intervention.

## Methods

### Sample collection

Sampling took place at the Arla Innovation Centre, in a pilot plant facility mimicking the industrial cheese making process for Herrgårdsost, located in Århus (Denmark), during 50 days between August and October 2019. Samples were taken from 3 replicate cheese production trials (named here PT1, PT2, PT3). The same source raw milk was used for all 3 replicates, but initially differential treatments: PT1 milk was pasteurised + microfiltered, PT2 was only pasteurised and PT3 was pasteurised + protein fortified. Pasteurisation was done at 72°C for 15 seconds. Microfiltration was performed using ceramic filters with a pore size of 1.2 µm (Tami Industries). For the protein fortification process, milk was ultrafiltered to 4.2 % target protein concentration, using a spiral membrane GR82 (Alfa Laval).

A total of 14 samples were taken for each trial, up to a total of 42 samples. Samples from the rind and core were taken during the ripening process.

Whenever cheese samples were taken from the rind of the cheese, they were taken ∼1 cm from the surface, while for the cheese core, they were taken at 2.5 cm deep from the surface, both on the same specific collection day for all 3 trials.

For each of the three batches, after the first ripening step (14 days at 10ºC), cheeses were split in 2 different lines, which were subject to two different ripening temperatures (13ºC and 20ºC) for a total of 35 additional days, taking samples after a total of 21 days and 42 days from the moment the milk was first processed. Samples had different consistency depending on the stage of the cheese process. The observed textures were: liquid milk, coagulated cheese grains (curds) and solid cheese. In order to obtain a liquid sample for further library processing from either the solid cheese or the coagulated cheese grains we liquified the samples using DNA/RNA Shield (Zymo Research) stabilization solution.

### DNA extraction and shotgun sequencing

DNA was extracted from 200µl volumes of each cheese sample, using the ZymoBIOMICS MagBead DNA kit (Zymo Research) following the manufacturer’s instructions, with the exception of excluding the last washing step, and final elution into 60µl of buffer EB (Qiagen). Subsequently, the DNA was fragmented using a Covaris LE220-plus Focussed-Ultrasonicator (Covaris), then converted into Illumina compatible shotgun sequencing libraries using the BEST protocol (Carøe, Gopalakrishnan, and Vinner, 2018.), prior to sequencing using 100 bp pair-ended chemistry on an Illumina NovaSeq6000 platform using Novogene’s commercial service.

### Bioinformatic analyses

#### Sequence data preprocessing

A total of ∼2,200 million paired end (Table S1) reads were generated (PT1: 863M, PT2: 658M,PT3: 682M). Read trimming and adapter removal was done using fastp (v. 0.23.4) (Chen et al. 2018), using the default Illumina adapters for trimming. Host DNA (*Bos taurus*; domestic cow) was removed from the reads by mapping the raw reads with bowtie2 (v. 2.5.0) (Langmead and Salzberg 2012) mapper against the cow reference genome (ARS-UCD1.3, *bosTau9*), and selecting the unmapped fraction with samtools (v. 1.19) (H. Li et al. 2009) (Table S1)

#### KHILL k-mers analyses

We used KHILL to do pairwise comparisons between each stage across all three samples. We randomly sampled 3 million reads from each stage and ran KHILL with default parameters. Since these comparisons are based on raw read libraries, we proceeded canonical k-mer counting without sketching, using all k-mer information contained in all 3 million reads.

#### Metagenomics analyses

We independently co-assembled each of the 3 sample batches (n = 14 × 3 = 42 samples) using the Anvi’o snakemake workflow for metagenomics (Eren et al. 2021; Köster and Rahmann 2012). The workflow runs several software steps which allow the user to go from raw FASTQ to a contig database and a profile containing status from the contig mapping step needed downstream to complete other analyses with Anvi’o. The steps followed by the workflow base quality of the reads is checked by the software illumina-utils v. 2.12 (Eren et al. 2013). MEGAHIT (v. 1.2.9) (D. Li et al. 2015) was used with Anvi’o metagenomic default parameters and only considered assembled those having a length >1kb. Mapper bowtie2 (v. 2.5.0) (Langmead and Salzberg 2012) was used to estimate coverage of contigs, by separately mapping back the FASTQs against the assembled set of contigs as reference sequences. At the end of the pipeline, a unique set of contigs (termed “contigs database”) and an Anvi’o profile with info like coverage and the tetranucleotide frequencies, was generated for each time point (Supplementary Data). The 42 profiles were merged, and together with the contig database constitute the starting point for downstream analyses and annotation with Anvi’o. Prodigal (v. 2.6.3) (Hyatt et al. 2010) was run once the contigs were assembled through Anvi’o to find genes by searching for ORFs, with Anvi’o default parameters. HMMER (v. 3.3.2) (Zhang and Wood 2003) was separately run through *anvi-run-hmms*, to identify and characterise genes from archaeal, protists, or bacterial origin, by comparing against single-copy core gene (SCGs) collections (Lee 2019), using hidden Markov models (HMMs). These collections of SCGs were used to get completion and redundancy of MAGs or bins. Additionally, ribosomal RNA HMMs were included in the search through barrnap (https://github.com/tseemann/barrnap), as included in the Anvi’o HMMs collection. A custom HMM profile for RNAPol-A/B was used as input as an extra parameter to the Anvi’o *anvi-run-hmms* command to specifically annotate contigs for these two genes.

Given the low complexity of our samples, manual binning was done for each group of samples, after running *anvi-interactive*. The Anvi’o interactive interface displays a central unrooted phylogenetic tree for the contigs, and GC content, coverage and taxonomic assignment are displayed. We considered a bin to be a MAG when completion was > 70%, and redundancy was < 5%. In the case of a tie, the longest bin was retained.

#### Taxonomy and functional analysis

To determine the taxonomy of our bins, we relied on Anvi’o collection of SCGs, which is based on Genome Taxonomy Database (GTDB) (Parks et al. 2018). *anvi-run-scg-taxonomy* program was run on the contigs database to annotate the SCGs present. Then, taxonomy assignment was done live during the manual binning process, by enabling the Anvi’o taxonomy option in the interactive interface, and exported afterwards to table format through the *anvi-summary* option. Kaiju (v.1.9.2) (Menzel, Ng, and Krogh 2016) was run with NCBI’s non-redundant protein database ‘nr’ (Pruitt, Tatusova, and Maglott 2005) as a reference to estimate taxonomy at the contig level. The Anvi’o tool *anvi-get-sequences-for-gene-calls* was first run on contigs to extract genes calls in fasta format, which were input for Kaiju. Output was next integrated into Anvi’o interactive visualisation interface, as described elsewhere: http://merenlab.org/2016/06/18/importing-taxonomy/ Krona (v. 2.8.1) (Ondov, Bergman, and Phillippy 2011) was used through function *kaiju2krona* from the Kaiju package over the Kaiju output, to convert the taxonomy into a hierarchical classification for the gene calls, for a manual overview of classified contigs. The interactive HTML Krona plots are available, for manual exploration, in the Additional Data section Classification of functions was done through Anvi’o (*anvi-run-ncbi-cogs*), by annotating the clusters of orthologous (COGs, version COG20 (Tatusov et al. 2000). A script included in Anvi’o (*anvi-compute-functional-enrichment-in-pan*) was run to calculate the functional enrichment in the pangenome, for the 143 *Lactococcus* genomes. COG assignment to major functional categories was done through a custom R script, by parsing the resulting file from the command above for the COG20 “functions” categories. Enrichments between *L. cremoris* and *L. lactis* strains were calculated as log2 fold change; this is, the log2 for the quotient ratio between *L. lactis* and *L. cremoris* assigned COGs (Supplementary Information).

#### Viral classification

Completion for viral MAGs was estimated by using CheckV (v. 0.9.0) (Nayfach et al. 2021), by running the *end_to_end* subroutine, using their native viral database v1.5.2. Dereplication for viral MAGs was done in two steps: i) we computed average nucleotide identity (ANI) for the viral genomes using pyANI (Pritchard et al. 2016), and ii) by using the Anvi’o function *anvi-dereplicate-genomes*, which returns the clusters and a representative MAG for each cluster. Standalone BLASTn algorithm as a command line interface (Altschul et al. 1990) was run against a non-redundant NCBI database, and the best hits for our viral sequences based on sequence length and similarity were sorted and filtered.

#### Relative abundances

To calculate relative abundances of each of our MAGs, we mapped back the raw FASTQ reads to our unique MAG catalogue. We started by indexing the MAG catalogue in FASTA format with bowtie2 (2.3.5) (Langmead and Salzberg 2012) and then reads were mapped with MAGs as a reference file with default parameters. As a result of the mapping step, SAM format files were transformed into BAM compressed files by samtools (v. 1.9) (H. Li et al. 2009) Next, using Anvi’o, we generated a contigs database for our unique MAG catalogue, and a merged profile for all these samples that were mapped to MAGs. We now generated an Anvi’o summary for each batch (PT111, PT2, PT3), reporting in a table the coverage for each MAG in the catalogue, for each batch. Using custom R (v. 3.5.0) (Team 2009) scripts with the coverage as an input, we extrapolated the approximate number of reads that were recruited by each sample, and next we used a compositional normalisation (R package *microbiome*; Lahti *et al*. 2017 http://microbiome.github.com/microbiome) to calculate the percentage of reads recruited by each MAG, which were plotted with R. The R package *phyloseqR* (McMurdie and Holmes 2013) was also used to handle relative abundances tables and facilitate plotting abundances.

#### Statistical analyses and plots

All plots done using the *ggplot2* package (Wickham 2011) and the *tidyverse* group of functions (Wickham et al. 2019) for R versions either v. 4.1 or v. 3.5 Basic two-way ANOVA analyses were performed with R (v. 4.1) (Team 2009). Package *effectsize* (Ben- Shachar, Lüdecke, and Makowski 2020) was used to estimate effect sizes and *pwr2* package was used to estimate statistical power for our design (https://cran.r-project.org/web/packages/pwr2/).

## Code availability

Code used to process the data can be found at: https://github.com/pollicipes/Metacheese. Additionally, data files generated after coassembly in Anvi’o CONTIGS.db format, together with FASTA files produced for each PT can be found at: https://sid.erda.dk/cgi-sid/ls.py?share_id=G0T7mMxBdg

## Raw data availability

Raw FASTQ sequencing data generated for the present study, together with corresponding metadata for each file, can be downloaded from: https://sid.erda.dk/cgi-sid/ls.py?share_id=C94UedvziI

## Acknowledgements

Danish Dairy Research Foundation award ‘Metacheese’ and DNRF143 award for funding.

## Author contributions

M.T.P.G, L.T.A, K.S and S.L designed and led the study, L.S.B, K.S, M.R and C.C carried out field work and provided samples. S.S.T.M, L.S.B, M.R and I.B.N extracted and prepared samples in laboratory, K.S, J.A.R performed computational analyses, A.N implemented KHILL method, T.S.P, L.T.A and M.T.P.G guided analyses, J.A.R, L.T.A and M.T.P.G interpreted the data, and J.A.R., L.T.A and M.T.P.G wrote the paper, with input from all other authors. All the authors approved the submitted version of the manuscript.

## Conflict of interest

None stated

## Notes

https://sid.erda.dk/cgi-sid/ls.py?share_id=C94UedvziI

https://sid.erda.dk/cgi-sid/ls.py?share_id=G0T7mMxBdg

### Competing Interest Statement

The authors have declared no competing interest.

