## Supplementary Information for "The effect of different milk pretreatment methods on microbiome community development during Herrgårds cheese production and ripening"

[5]*Arla A/S*

[6]*University Museum, NTNU, Trondheim, Norway*

[#]*Equal contribution*

### *Lactococcus* MAGs specific analyses

##### *Lactococcus* genome analyses

To confirm our potential observation of exactly 2 strains of *Lactococcus*, we annotated the contigs for both RNA polymerase A and B (RNAPolA/B), using the anvi'o function *anvi-run-hmm* to annotate our 3 contigs database (PT1, PT2 and PT3). This program relies on Hidden Markov Models (HMM) to fastly identify sequences in genomes (Zhang and Wood 2003). RNAPolA/B is a SCG and each bacterial genome should carry exactly one copy of each (Gaïa et al. 2023). As expected, we identified exactly one copy of each for each *Lactococcus* assigned bin (Table S1).

Genome completion estimates for either *Lactococcus lactis* strains in each batch were between 52-69%, despite genome length being for all of them between 1.8-2.0 Mb. These sizes align well with complete reference genomes already published for this species. One of our binned *Lactococcus* strains had consistently higher coverage across all the 14 processing steps for the 3 batches (average 9.2 to 15.5x more than the other strain). Given the observed lengths being within the expected range we believe the lower completion reported by anvi'o is an artifact of the differential coverage between strains. As we said, the coverage is on average homogeneous across each strain, except for a few SCGs, which may be higher for the low coverage strain, and hence why these SCGs are assigned by anvi'o to the strain with more coverage (this is; higher abundance). Given that the completion is estimated through the presence in bins of annotated SCGs, anvi'o missinterprets and underestimates completion of the genome. Indeed, if we do the binning alternatively by not attending to these coverage issues we can get up to a 100 % completion for one of the *Lactococcus* genomes, while for the other one we still get < 60%. Although, in this last case, we will be assembling a sequence "chimera" for the complete genome. In light of this, and to be on the safer conservative side, we proceeded to bin only those homogeneous coverage contigs through the interactive interface. This guarantees that we have a correct genome length, while saving differences between the species, despite underestimation of completion. A similar issue has been already reported in (Delmont et al. 2018). Furthermore, an interactive and hierarchical view of the Kaiju contig-level taxonomy can be explored through the Krona (Ondov, Bergman, and Phillippy 2011) interface (Additional Dataset)

##### *Comparative phylogenomics identifies Lactococcus spp. strains*

We hypothesize that both *Lactococcus* genomes we recovered are derived from the cultures added at the inoculation step (Fig 1A; main text). As we were not successful in recovering a full clean genome in a bin, we will build a phylogenetic tree incorporating other *Lactococcus* reference genomes. We downloaded 100 NCBI complete reference genomes for 12 different *Lactococcus lactis* subspecies (*carnosus, cremoris, garvieae, lactis, paracarnosus, petauri, piscium, protaetiae, raffinolactis, taiwanensis*; plus 4 undetermined *Lactococcus* genomes, also available in the database). Furthermore, we added the 37 starter culture *Lactococcus* genome sequences available to us, which had been already previously sequenced (Lene Tranberg, personal communication; Additional Data, Tables S6). In total, summed up with our 6 *Lactococcus* bins (2 x PT), we had a total of 143 complete lactococcal genomes. Genome completion for the 137 additional external genomes was assessed with anvi'o and all of them were ≥ 93% complete, with < 3% of redundancy (Table S6).
To identify the single-copy core genes (SCG) (Lee 2019), that could be informative to build a phylogenetic tree and classify our sequences, we computed a pangenome with our 143 Lactococcus genomes using anvi'o pangenome tools (Methods). Results revealed that our 6 bins clustered equally 3 & 3 into two differentiated groups, together with either all of *Lactococcus lactis subsp. lactis* or *Lactococcus lactis subsp. cremoris*. A set of 223 SCG clusters across all 143 *Lactococcus* genomes + bins were selected. An unrooted maximum likelihood phylogenetic was constructed with the 223 SCG clusters (Figure S6). The resulting tree indicates a clean separation of both *cremoris* and *lactis* strain groups, evenly assigning our 6 lactococcal bins to either group. Upon examination of the tree, we were able to visually pinpoint the closest culture strain partially recovered in our bins. For the *cremoris* strain the closest candidate was CRE6_0 and LAC2_0 for *lactis* (Figure S6. This is useful to substitute now our candidate *Lactococcus* bins with the corresponding exact culture strain, being more complete

than our binned lactococcal partial genomes. All other *Lactococcus* external genomes also clustered together with their relatives, and the unspecified *Lactococcus* genomes in NCBI were closely related to *garvieae* subsp.

***Lactococcus pangenome***

We built a pangenome for the 143 *Lactococcus* genomes using anvi'o pangenome commands, as instructed in https://merenlab.org/2016/11/08/pangenomics-v2/. In the interactive display of pangenome we selected the SCG clusters, common to all the 143 strains of *Lactococcus* used. Metadata including the strain common name, subspecies, origin and code for the 143 genomes was manually curated and added to the pangenome for improved viewing. MCL clustering algorithm (van Dongen and Abreu-Goodger 2012) was used through anvi'o to identify gene clusters, using default parameters.

***Lactococcus phylogeny***

A total of 223 SCG gene clusters were used to build a phylogenetic tree. This was used to build a maximum likelihood phylogenetic tree with FastTree2 (Price, Dehal, and Arkin 2010), as implemented in anvi'o. A Newick format tree was obtained and viewed alongside metadata through the anvi'o interactive interface.

***The lactis subsp. is enriched in higher energy-related metabolic processes.***

Next, to quantify metabolic capacities of both strains that could be affecting cheese final characteristics, or that may help explain patterns of abundance later on, we annotated the genomes of both *lactis* (n = 52) and *cremoris* (n = 54) with the clusters of orthologous genes database (COG20) (Tatusov et al. 2000) , using anvi'o function to annotate COGs. This provides a functional annotation for the gene clusters. Then, using the custom anvi'o function *anvi-compute-functional-enrichment-in-pan* we identified the functions significantly overrepresented in either *lactis* or *cremoris* versus any other *Lactococcus* species genome, that is; those exclusive to any of our two strains in question, and selected those functions significant at p-value < 0.01 (Figure S6). COG terms can be further classified at a higher level by categorizing the functions within a classification represented with the 26 alphabet letters, from A to Z, each one representing a broad function category. Thus, next, we classified each group of our significant functional terms for each of the two strains into this systematics and calculated the enrichment at the level of categories by comparing the log2 fold-change of *lactis* functional categories over *cremoris* functions.

In general, functions regarding energy-related processes like inorganic ion transport (logFC: 2.46x), lipid transport (logFC: 1.58x), coenzyme transport (logFC: 1.00x) and carbohydrates transport (logFC: 0.78x) are prevalent functions in *lactis* strains, together with more general unclassified energy production and conversion processes, which are also enriched (logFC: 0.74x). The presence of citrate-related enzymes (citrate lyase synthetase, CitC, and its alpha subunit) in *lactis*, matches the subsp. *lactis* biovar diacetylactis, (known to be present in our cultures), which gives the cheese a distinct odor, after converting citrate into diacetyl, a strong odorous compound (Fernández et al. 2011). Other processes such as defense mechanisms or secondary metabolites biosynthesis are more prevalent in *cremoris* strains (Figure S6). Incidentally, within the inorganic ion transport category, in subsp. *lactis* we detected a gene encoding a key iron binding protein (Fra/YdhG), from the frataxin (Fra) family, which transfers iron to the biosynthetic cluster of iron-sulfur, acting as a key iron channeling molecule (Bencze et al. 2006). Unless supplemented, cheese is a highly iron-limited environment, as milk is naturally poor in iron, while containing lactoferrin, which is an antibacterial compound (Monnet et al. 2015). Those bacteria being capable of acquiring iron more efficiently, may thrive better than others, as iron is a growth-limiting environmental factor in complex cheese rind microbiomes (Saak et al. 2023; Monnet et al. 2015). Taken together, the iron-uptake related improved capacities and the specific energetic transport functions in subsp. *lactis* may contribute to a higher abundance than *cremoris* strain.

### Supplementary Figures

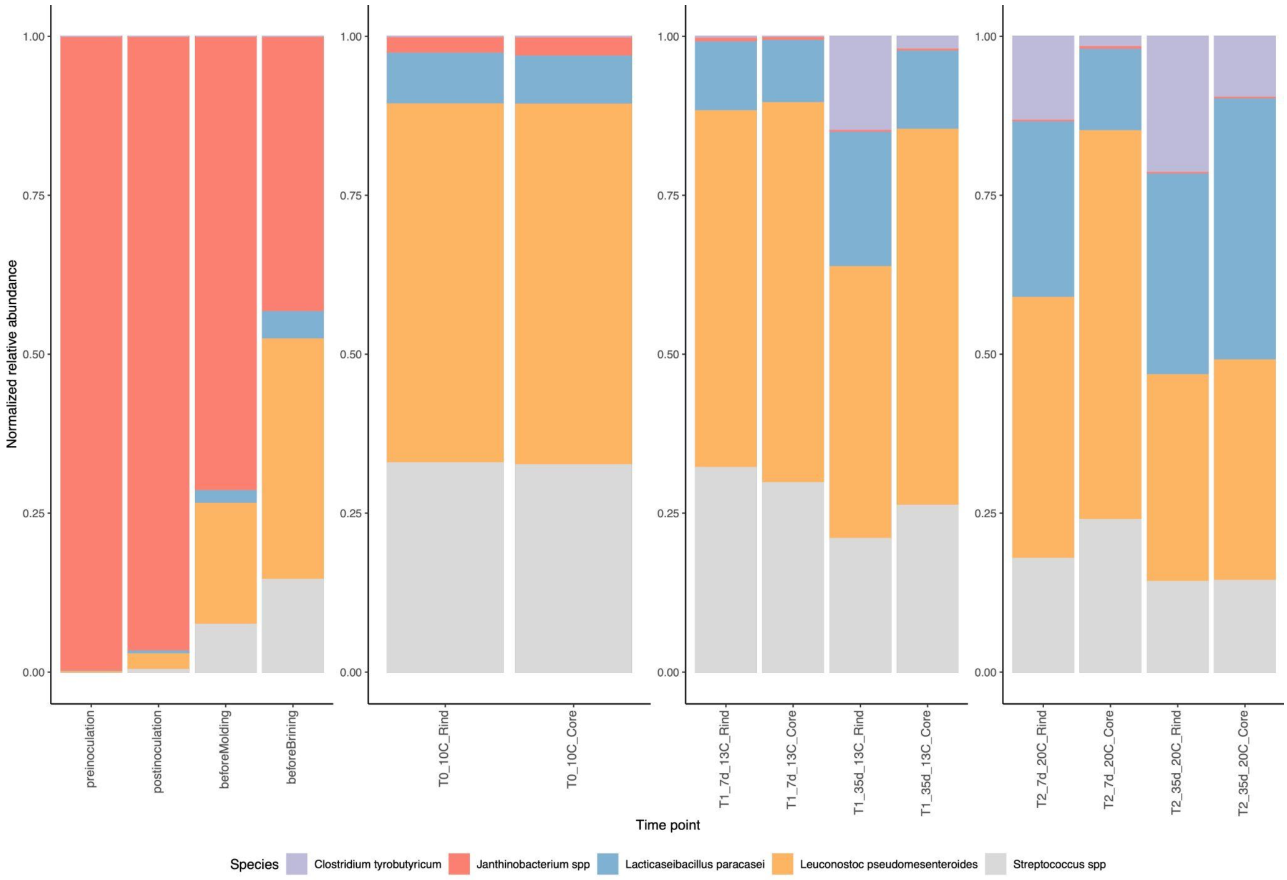


**Figure S1:** Abundance of minor abundant bacteria.

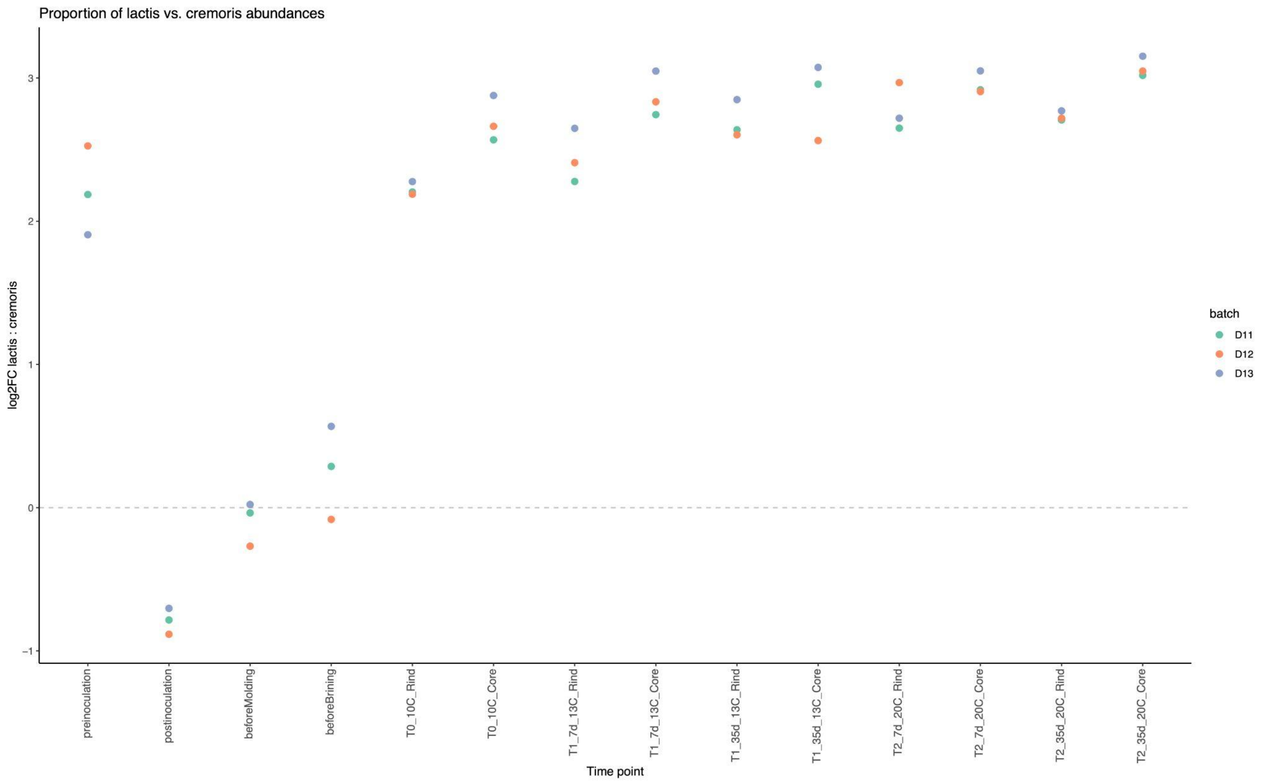


**Figure S2:** Proportion *Lactococcus lactis:Lactococcus cremoris* across the whole process for the 3 PTs

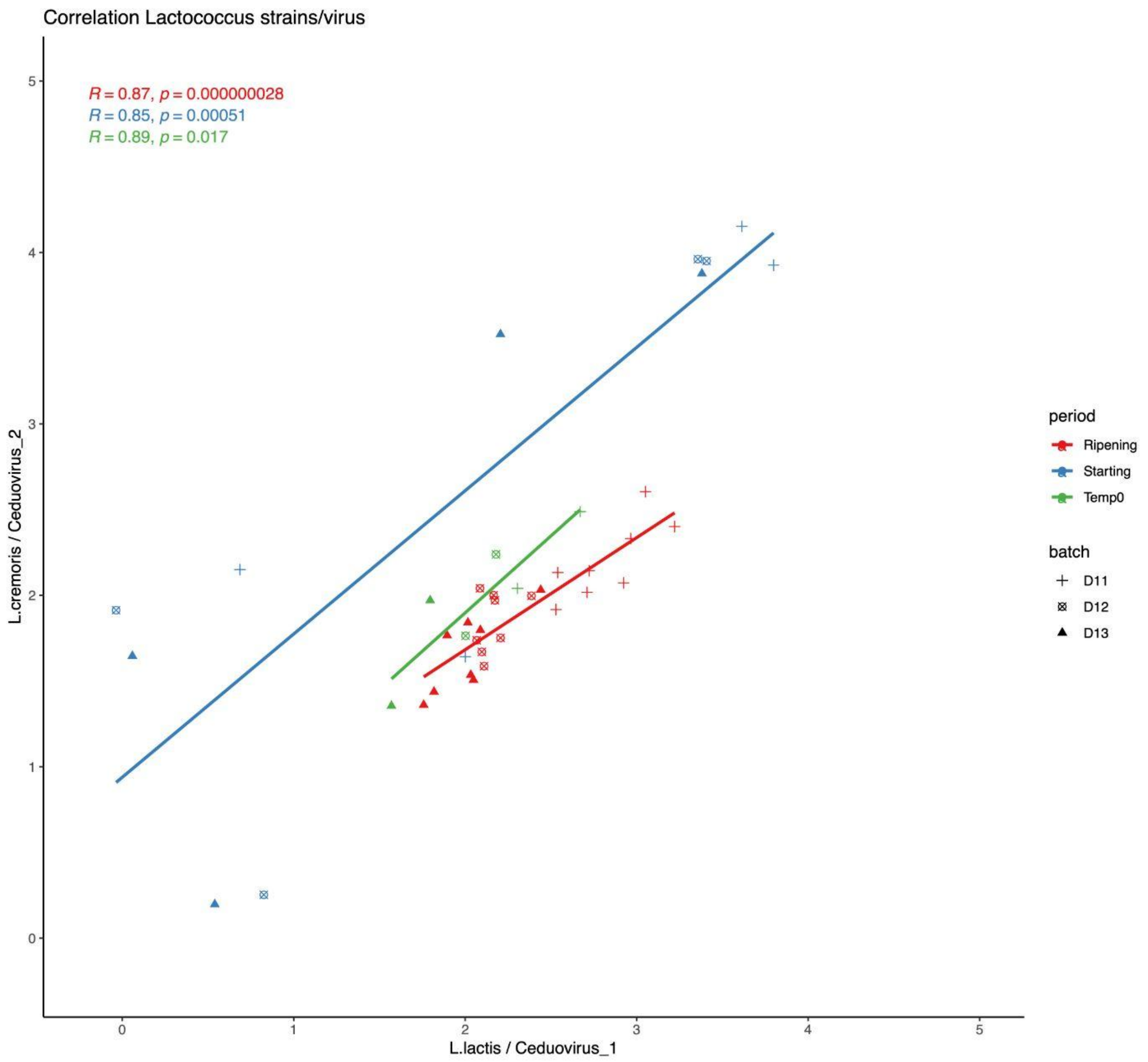


**Figure S3:** Correlation Lactococcus strains/virus. X-axis shows how the proportion between the *L. lactis lactis* and its associated phage is constant across the whole process, and this ratio of proportion maintains proportional to the ratio of *L. lactis cremoris*. relative to its phage associated, despite the changes in the proportions for both bacterial strains.

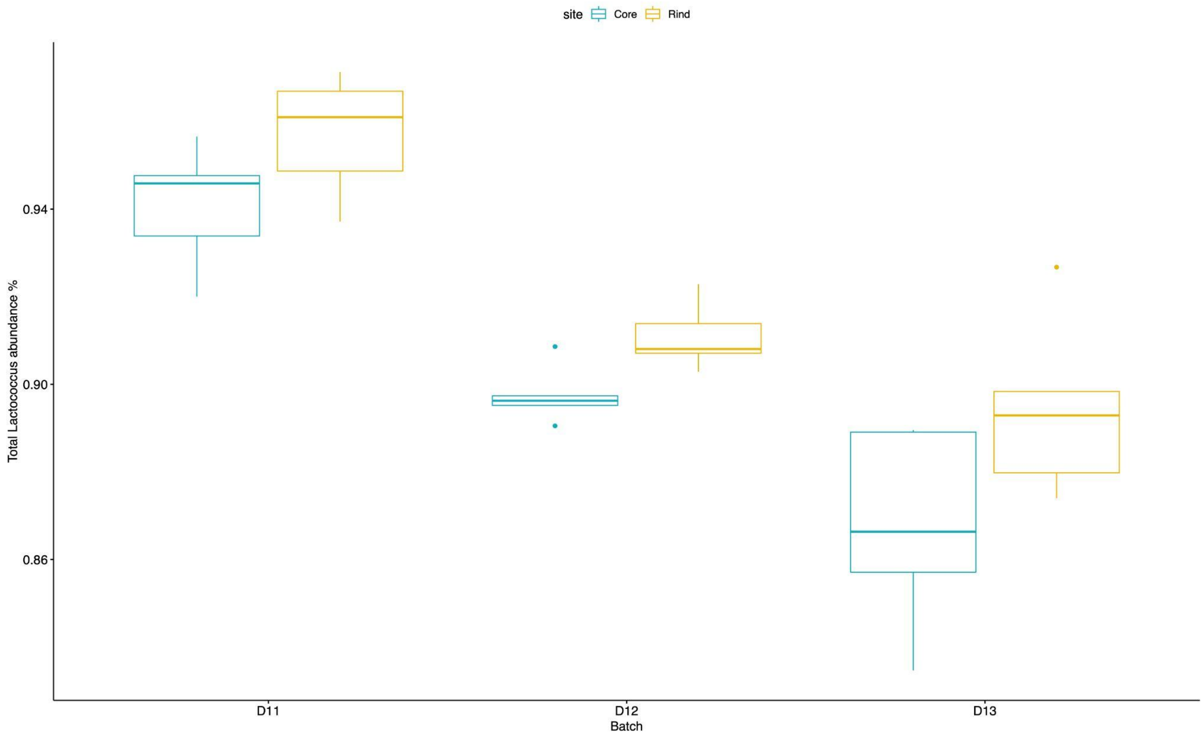


**Figure S4:** Lactococcus abundances by rind/core.

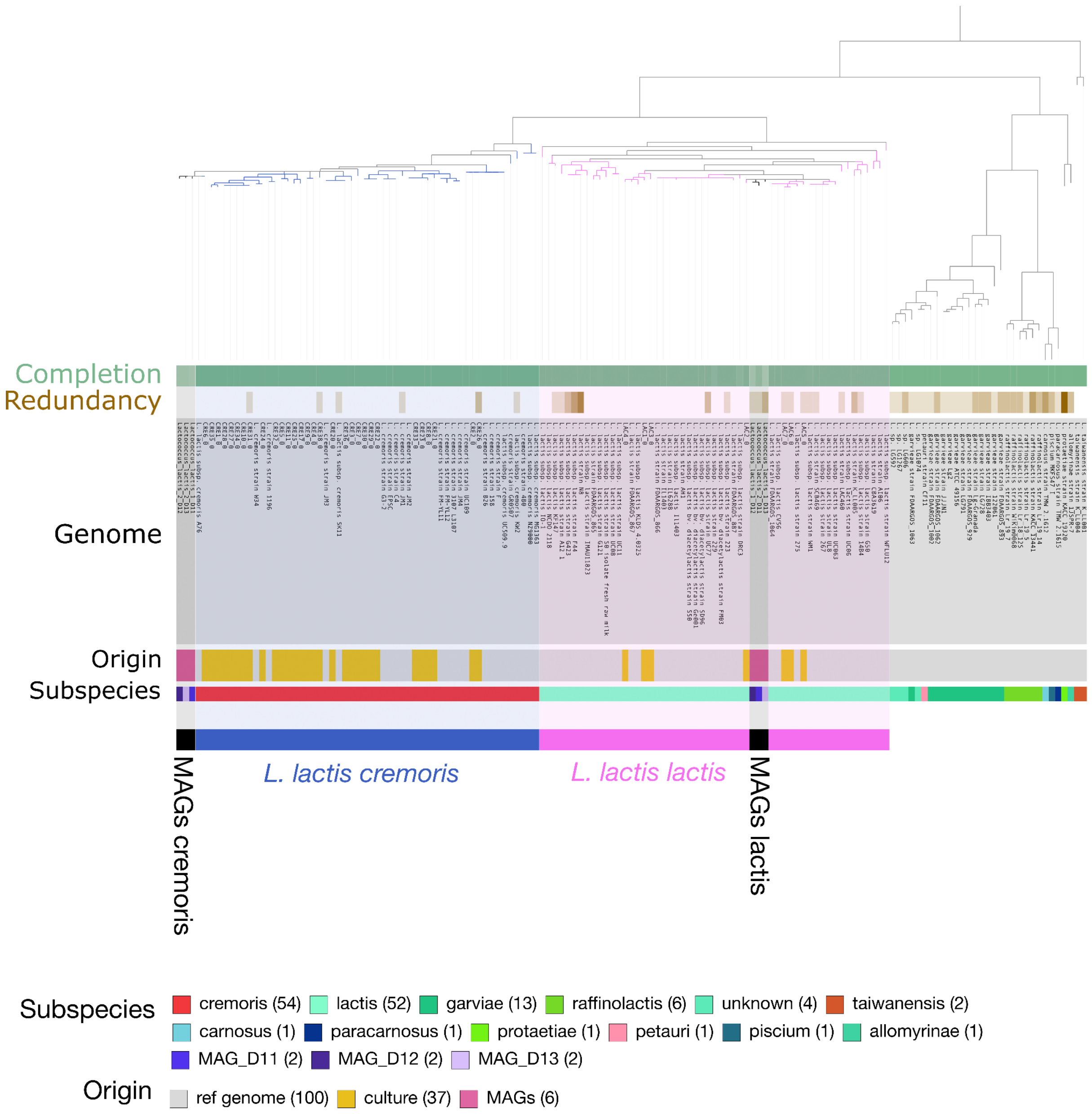


**Figure S5:** *Lactococcus* genus phylogeny for the 143 genomes, based on 223 SCG.

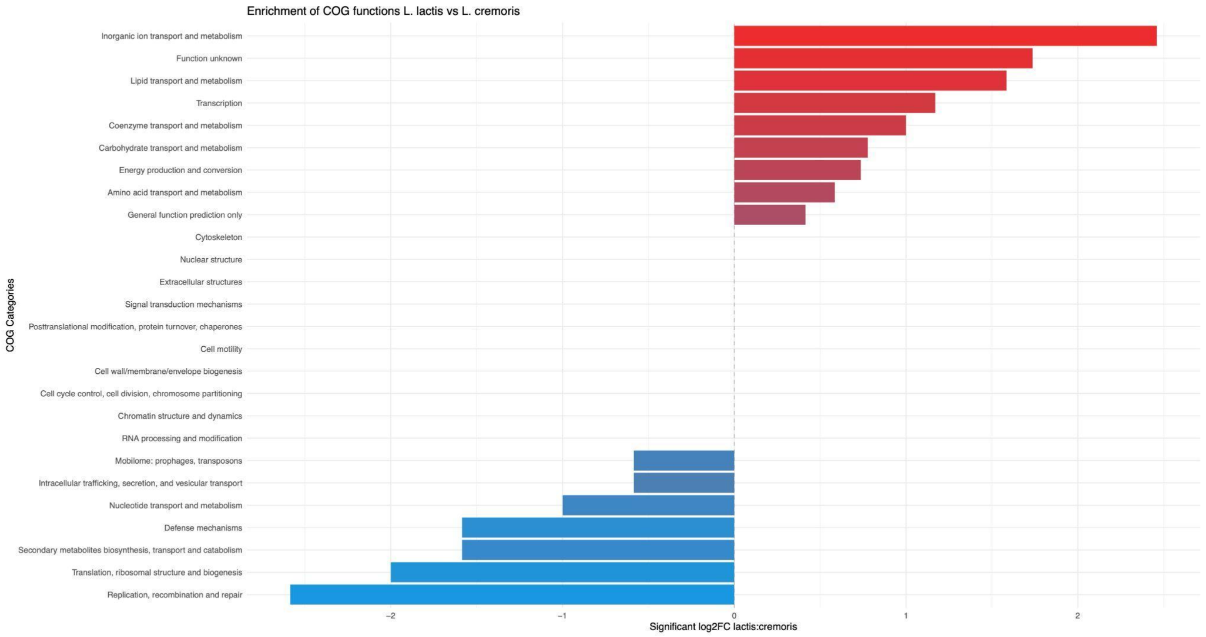


**Figure S6:** Enrichment of COG20 functions between lactis and cremoris strains. Red scale bars represent broad categories for COGs enriched in lactis, while blue tone bars represent those enriched for cremoris.

| | **RIPENING STAGES** | | |
|---|---|---|---|
| | **PT1** | **PT2** | **PT3** |
| **Average Reads** | 238999 | 232135 | 274328 |
| **SD** | 95036 | 88571 | 127381 |
| **Average % abundance** | 0.2 | 0.27 | 0.28 |

**Table S10:** Average number and percentage of reads mapping to MAGs other than *Lactococcus spp.* (n = 2 MAGs) or bacteriophages (n = 2 MAGs) during ripening stages (steps 5-9 in Table 1)
